# Rationales for microbial conservation: A quantitative global assessment

**DOI:** 10.64898/2026.02.02.703404

**Authors:** Robert R. Junker

## Abstract

Despite the foundational role microorganisms play in sustaining life on Earth, they have been largely overlooked in global conservation agendas, driving the emergence of microbial conservation as a critical discipline. While major assessment reports successfully mobilize support for the conservation of macroscopic biodiversity by documenting its value, threats, and intervention effectiveness, comparable evidence for microbes is lacking. I provide this missing evidence by synthesizing 33,297 effect sizes across three second-order meta-analyses. These analyses (1) identified land-use and land-cover change as well as specific pollutants as the primary threats to microbial diversity, function, and community integrity, (2) demonstrated the essential ecosystem services microbes provide, and (3) revealed the insufficient microbial conservation gain achieved by existing interventions. Building on these insights, I revisit the concept of vulnerability to propose targeted microbial conservation strategies that maintain or restore microbial diversity and function. The evidence presented here underscores the urgency of integrating microbes into nature conservation, thereby protecting the very foundation of life and safeguarding ecosystem integrity and planetary health.

## Introduction

Conservation interventions can effectively counteract the ongoing decline of biodiversity, habitat loss, and ecosystem degradation (Brondízio et al. 2019; Langhammer et al. 2024). As conservation competes with other global crises for attention and resources (Dasgupta 2021), rigorous evidence is essential. To justify investment, we require clear data on ecosystem functions, threats, and the efficacy of specific conservation actions. Such evidence is widely synthesized in national and international reports (Brondízio et al. 2019; Wirth et al. 2024; Secretariat of the Convention on Biological Diversity 2020) that document the instrumental value of plants and animals (and some macro-fungi), identify drivers of biodiversity loss, and evaluate conservation measures for these taxa and their habitats.

Yet, the virtual absence of two entire domains of life (Bacteria and Archaea) and eukaryotic microorganisms from these assessments is increasingly recognized, positioning microbial conservation as the next frontier in safeguarding biodiversity and ecosystem function (Gilbert et al. 2025; Gilbert et al. 2025; Junker and Farwig 2025). Ranging from the production of fermented foods like chocolate (Gopaulchan et al. 2025) to the mitigation of climate change (Peixoto et al. 2024), the instrumental value of microbes across all scales of life is indisputable. Conservation action becomes imperative when anthropogenic global change threatens microbial diversity and composition, undermining the foundation of their functions and services. Current evidence that microbial diversity and function are at risk comes primarily from laboratory experiments (Rodríguez Del Río et al. 2025) or meta-analyses focusing on individual drivers (Sun et al. 2025). To develop targeted microbial conservation strategies, we require comprehensive assessments of the drivers threatening microbial diversity and function. Identifying primary threats will enable the design of effective interventions to protect microbial communities and their functional associations with ecosystems and hosts. Crucially, evaluating how existing macro-organism conservation affects microbes will help distinguish where current measures suffice and where dedicated microbial strategies are needed.

To establish an empirical basis for microbial conservation, I performed three second-order meta-analyses (SOMA). These analyses aim to: (1) identify and rank the global change drivers threatening microbial diversity, composition, and function; (2) quantify the instrumental value of microbes for ecosystem services and host health; and (3) determine the impact of existing conservation interventions on these community properties. Synthesizing the results of meta-analyses provides a high level of scientific evidence, as this approach incorporates thousands of effect sizes to enable conclusions based on the full body of available data (Makowski et al. 2023). The specific drivers, functions, and interventions analyzed here were defined by a systematic literature search rather than a priori selection, capturing the themes most extensively studied by the scientific community and thereby reflecting the prevailing research priorities in microbial ecology. Drawing on this evidence, I adapted the established vulnerability concept, composed of exposure, sensitivity and adaptive capacity (Turner et al. 2003; Foden et al. 2013) of microorganism to global change, to identify microbial systems most at risk from global change.

Integrating microbes into current conservation strategies offers immediate benefits to established conservation goals (Junker and Farwig 2025), from increasing host health (Peixoto et al. 2022; Trevelline et al. 2019) (species protection) to enhancing habitat restoration (Contos et al. 2025; Hart et al. 2020) (habitat protection). However, defining explicit targets to protect microbial diversity and function marks a critical next step in the field, acknowledging that these invisible constituents provide or support virtually all ecosystem services (Gilbert et al. 2025; Gilbert et al. 2025; Junker and Farwig 2025). This synthesis lays the empirical foundation to establish microbial conservation as both a coherent discipline and a practical field of application, advancing an integrated framework that accounts for the functions of all domains of life.

## Methods

### Systematic literature search and selection criteria

In September 2025, I searched the web of science (advanced search) for meta-analyses on (1) global change drivers that affect microbial communities in terms of abundance, biomass, diversity, composition, and functionality; (2) the instrumental values of microbes in supporting ecosystem services and host performance; and (3) microbial responses to existing conservation interventions. Based on the literature, I conduced three second-order meta-analyses (SOMAs). Such analyses that summarize the results of meta-analyses are regarded as one of the highest levels of scientific synthesis, as this approach can incorporate thousands of effect sizes and thus enables conclusions based on the full body of available evidence (Makowski et al. 2023). Full methods and the search terms for the SOMAs are listed in SI.

I exported full records of the results of the searches as Tab Delimited Files and used the R package *metagear* (Lajeunesse 2016) to screen candidate studies following PRISMA (Preferred Reporting Items for Systematic Reviews and Meta-Analyses) recommendations. I screened the title and abstract of each article to assess its eligibility for inclusion in the final analysis. The main criteria required a quantitative meta-analytical assessment of studies that reported results in the context of the three intended SOMAs. I excluded meta-analyses in the medical or veterinary context. The resulting initial set of articles was then downloaded and evaluated based on the following criteria: Only studies on microbial assemblages, bacteria, or fungi were included; studies focusing on a small taxonomic range (e.g. single strains) were excluded; only studies reporting log response ratios (LnRR) or effect size measures that can be readily transformed into LnRR were included (Fox 2022), thus studies reporting measures based on mean differences (e.g. Hedges *d*) were excluded as they cannot be transformed into response ratios (Tamburini et al. 2020). Some meta-analyses reported main effects alongside additional effect sizes addressing study-specific questions. In these cases, only the main effects were extracted to ensure data independence. The selection of ecosystem functions, drivers, and interventions considered in the three SOMAs was not pre-defined but emerged directly from the systematic search strategy; consequently, the analyzed categories reflect the current focus of the scientific community and represent the most extensively studied, and thus arguably the most critical themes in microbial ecology. The selection steps and resulting study numbers, including exclusions, are shown in PRISMA flow charts in SI (Fig. SI.1). From the final selection of published meta-analyses, I extracted mean effect sizes (log response ratios, LnRR) and standard errors *SE* from tables, supporting information or from figures using the WebPlotDigitizer 5.2 (automeris.io) (Rohatgi, n.d.; Jelicic Kadic et al. 2016). A full list of meta-analyses included in this study is given in SI and can be found here: https://doi.org/10.6084/m9.figshare.30875873.

For the SOMA on threat-based rationales, each effect size was assigned to one global change driver category following the IPBES (Brondízio et al. 2019) framework (climate change, exploitation, invasive species, land use, pollution). However, to account for the fact that microorganisms and microbial communities respond to specific physicochemical cues rather than broad conceptual categories, these broad drivers were further stratified into 15 specific factors (see Fig. 1B). Pooling distinct and opposing factors (e.g. drought and increased precipitation within ‘climate change’) masks significant biological signals and inflates unexplained heterogeneity. The factors elevated CO₂, drought, fire, increased precipitation, and warming were assigned to ‘climate change’. Fire was included in this category as a disturbance factor increasingly driven by climatic shifts (e.g., wildfire regimes), although I acknowledge it can also be classified as ‘land use’ when used for management (prescribed burning). Since the underlying meta-analyses included both fire types, I tested the robustness of this classification; reassigning fire to ‘land use’ had negligible impact on the calculated effect sizes (Fig. 1B) and did not alter the overall interpretation of results. I assigned heavy metals, microplastics, ozone, and nutrient enrichment (nitrogen and phosphorus addition) to the category ‘pollution’. Although N and P addition are frequently linked to agricultural intensification and could theoretically be categorized under ‘land use’, I classified them as pollution to reflect their mechanistic role as chemical alterations of the environment. This distinction aligns with the IPBES assignment and separates these chemical drivers from the physical habitat disturbances (e.g., tillage, mowing) characteristic of the ‘land use’ category. Accordingly, I assigned mowing and various land-use and land-cover change (LULCC) factors, specifically land conversion, general disturbances, forest degradation, conversion of forest to agriculture, and understorey removal, to the category ‘land use’. Although mowing represents a recurrent management practice (intensification) rather than a permanent change in land cover, I included it here because, like degradation and conversion, it acts as a physical driver altering habitat structure and biomass. This classification effectively separates these physical habitat modifications from the chemical drivers categorized under ‘pollution’ and ‘climate change’.

**Fig. 1.**
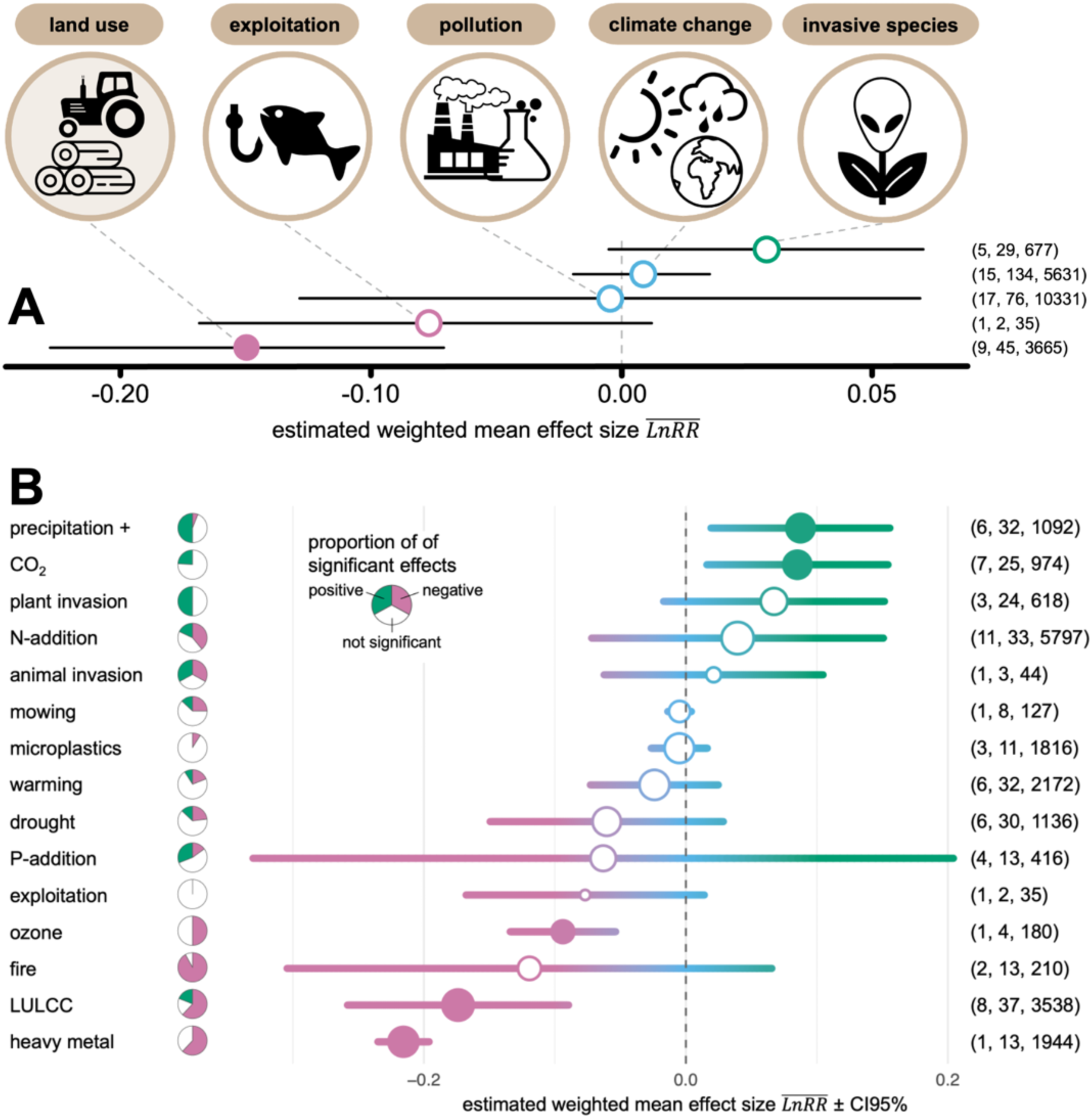
Impacts of global change drivers and factors on microbial diversity, abundance, biomass, and functionality. Mean effect sizes (± 95% CI) of (**A**) broad global change drivers and (**B**) further stratified specific factors. Filled circles denote significant effects. Color gradients visualize the heterogeneity of responses within categories ranging from negative (pink) to positive (green) with rather neutral effects in blue. Circle size is proportional to the number of original effect sizes. Pie charts inform about the proportion of positively (green) or negatively (pink) significant effects as reported in original meta-analyses. Parentheses denote sample sizes of meta-analyses, extracted estimates, and of underlying primary effect sizes. LULCC = land-use and land-cover change.

Although some effect sizes originate from single meta-analysis, these sources were retained because they synthesize a substantial volume of primary research (large number of effect sizes and primary studies). Excluding these comprehensive assessments would disregard the most robust available evidence for those specific factors. Therefore, statistical weight and reliability were interpreted based on the underlying cumulative sample size rather than the count of meta-analyses alone.

SOMAs may violate the assumption of independence among effect-size estimates when primary meta-analyses draw on overlapping datasets. To assess this potential bias, I downloaded the lists of studies used in the primary meta-analyses (where available) and checked for overlaps among those reporting on similar effects (driver × response combinations) and thus contributing to the same effect-size estimates shown in Fig. 1–3 of the main text. Overall, overlaps were detected in only four primary meta-analyses (indicated with an asterisk * in the reference list in SI), with shared studies accounting for 2.6%–15.3% (mean 9.3%) of their datasets. Therefore, any potential bias due to shared datasets is limited to few primary meta-analyses and, where present, affects only a small proportion of studies. These low overlap levels suggest that the overall results are not meaningfully affected.

**Fig. 2.**
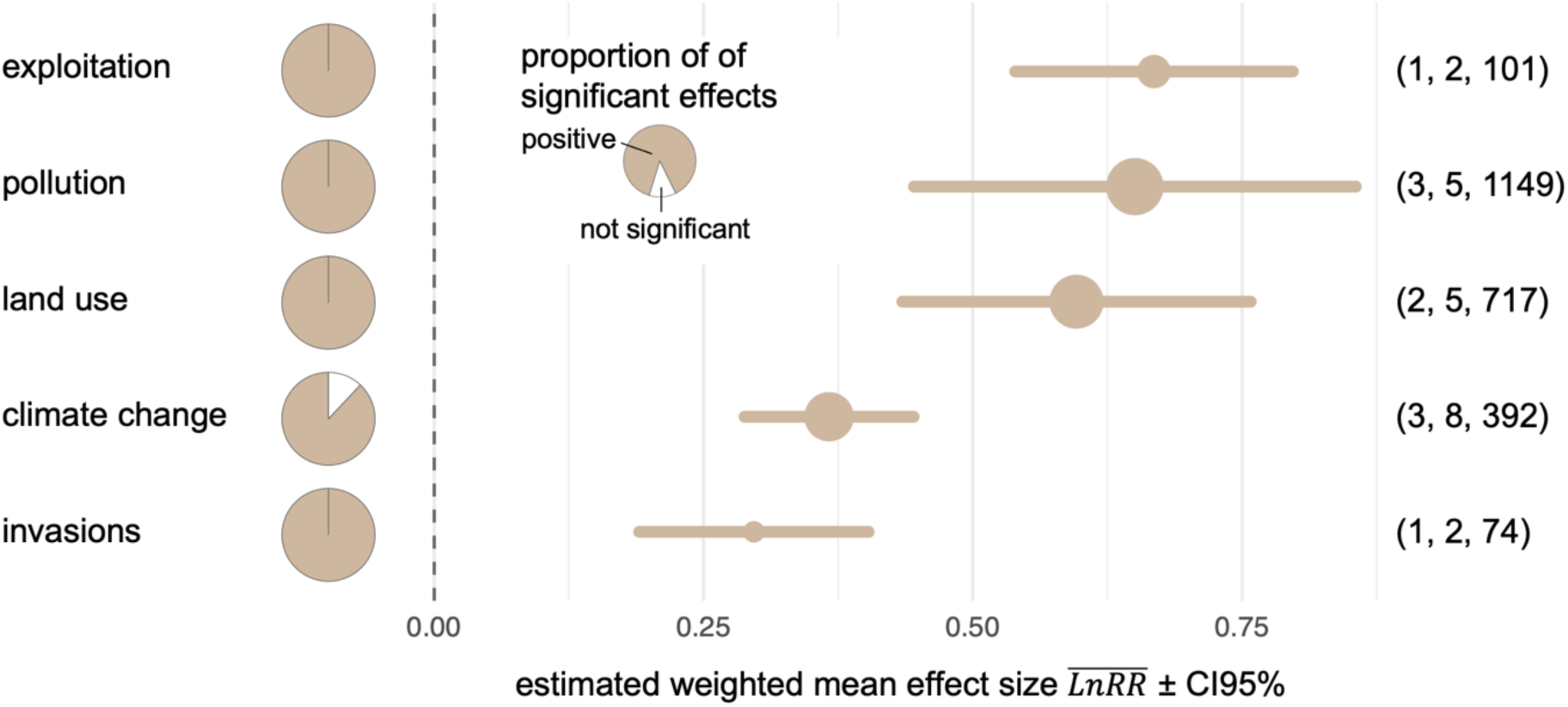
Impacts of global change drivers and factors on microbial community composition. Effect sizes ± CI95% of global change drivers. All effects are significant. Circle size is proportional to the number of original effect sizes. Pie charts inform about the proportion of significant effects as reported in original meta-analyses. Parentheses denote sample sizes of meta-analyses, extracted estimates, and of underlying primary effect sizes.

**Fig. 3.**
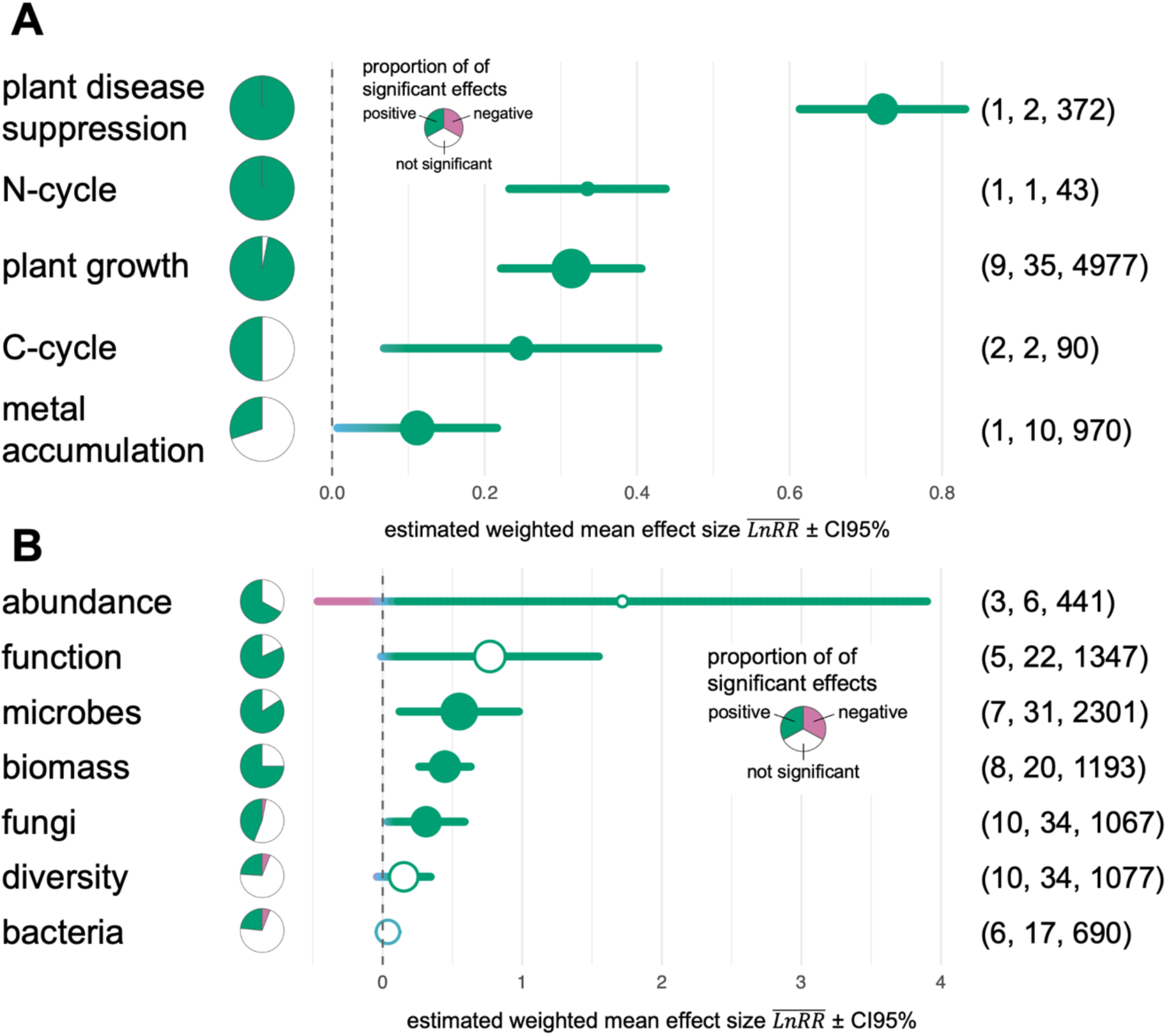
Microbial effects on ecosystem functions and host performance (A) and effects of existing conservation interventions on microbial abundance, biomass, diversity, function, and taxa (B). Effect sizes ± CI95% are shown. Filled circles denote significant effects. Color gradients visualize the heterogeneity of responses within categories ranging from negative (pink) to positive (green) with rather neutral effects in blue. Circle size is proportional to the number of original effect sizes. Pie charts inform about the proportion of positively (green) or negatively (pink) significant effects as reported in original meta-analyses. Parentheses denote sample sizes of meta-analyses, extracted estimates, and of underlying primary effect sizes.

### Statistical analysis

All statistical analyses where performed using the statistical software R (R Core Team 2025). Based on the resulting dataset, I ran mixed-effects meta-analyses with *SE^2^* as sampling variance using the function *rma.mv* implemented in the R package *metafor* (Viechtbauer 2010) that allows to define fixed effects (moderators) and random effects that control for non-independence in the data due to multiple effect sizes obtained from most meta-analyses. The global model was defined as follows:

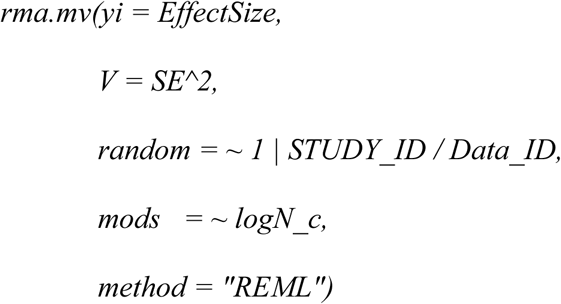

with *EffectSize* = LnRR, *SE^2* = sampling variance obtained from original meta-analyses, the *random* effect to control for non-independence in the data due to multiple effect sizes obtained from most meta-analyses, *logN_c* = centered log of the number of studies included to estimate effect size in original meta-analyses to test for sample size bias, and *method* = *Restricted Maximum Likelihood*. To evaluate whether sample size explained additional heterogeneity among effect sizes, I compared a full multilevel model including the moderator (*logN_c*) with a reduced model lacking this term using a refitted likelihood-ratio test, ensuring valid comparison of nested models. No evidence of sample size bias was detected; therefore, models without this moderator were used in subsequent analyses. For each analysis, I extracted the number of meta-analyses considered, the number of samples, the number of original effect sizes as well as the number of meta-analyses that reported either positive or negative effects. In cases where results are based on a single meta-analysis, I ran fixed effects models excluding the random factor:

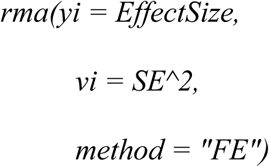

For N-cycle (Fig. 1B) only one effect size was available, in that case I used the effect size and SE from the original meta-analysis. To test whether global change drivers, microbial taxa, or α-components of microbial communities (abundance, biomass, diversity, functionality) differed in their mean effects, I fitted multivariate mixed-effects meta-analytic models with the relevant moderator. Moderator significance was evaluated using the QM statistic, which tests whether categories show distinct mean effects.

To assess potential publication bias, I performed Egger’s regression test by regressing standardized effect sizes (EffectSize/SE) on their precision (1/SE), using STUDY_ID-cluster-robust standard errors. In cases where only a single study contributed to the meta-analytic model, robust estimation was not possible, and standard Egger regression results were obtained using ordinary least-squares summaries. Additionally, I visually inspected funnel plots for evidence of small-study effects, such as asymmetry, uneven dispersion, or missing studies on one side of the plot.

## Results and discussion

### Threat-based rationales

To summarize evidence of threats posed to microbial diversity, abundance, biomass, and functionality, a systematic search was conducted in Web of Science for meta-analyses addressing microbial (bacteria, fungi, the total microbial assemblage) responses to factors of global change. Effect sizes were assigned to broad global change driver categories commonly used by frameworks like IPBES (Brondízio et al. 2019) (Fig. 1A). These were further stratified into 15 distinct factors (Fig. 1B) to account for factors with opposing physicochemical effects within the broad categories (e.g., drought *versus* increased precipitation within ‘climate change’). The meta-analyses reported either on global change effects on *⍺*-scale metrics (defined as the inventory or state of a single unit, including abundance, biomass, diversity, and functionality) or on shifts in community composition (*β*-scale metrics).

The overall effect of global change drivers on *⍺*-scale metrics of microbial communities showed a non-significant negative trend (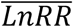 = −0.03; 95 % *CI* = −0.077 to 0.018; *p* = 0.221, based on *n* = 20,339 effect sizes). Consistent with this lack of a unified signal, broad global change driver categories varied significantly in their mean effects on α-scale metrics of microbial communities (Fig. 1A, *QM_4_* = 12.67, *p* = 0.013). Land use emerged as the strongest driver of declines in microbial *⍺*-scale metrics. While other drivers did not significantly deviate from zero, their large 95% confidence intervals indicate pronounced heterogeneity of effects within these broad categories (Fig. 1A). Reflecting this heterogeneity, the 15 specific global change factors explained a considerable proportion of the variation in α-scale metrics (Fig. 1B, *QM_14_* = 61.68, *p* < 0.0001). While increases in precipitation and CO_2_ concentration on average significantly increased *⍺*-scale metrics of microbial communities by 9.1 and 8.9 %, increases in ozone concentration, land-use and land-cover change (LULCC), and heavy metal pollution significantly decrease *⍺* α-scale metrics by −9.0 %, −16.0 %, and −19.4 %, respectively. Of the 15 global change factors investigated, eleven exhibited negative mean effect sizes, yet only three deviated significantly from zero. This lack of significance reflects substantial heterogeneity, i.e. large 95% confidence intervals (Fig. 1B). Likewise, out of five global change drivers with positive mean effect sizes only two differed significantly from zero (Fig. 1B). Considering the proportion of significantly positive or negative effect sizes within factors in the original meta-analyses further highlights the strong variability in global change effects and presumably context-dependency (Fig. 1B). While effect sizes did not differ significantly between microbial taxa (*QM_2_* = 1.690, *p* = 0.430), suggesting broad taxon-independence, the specific types of *⍺*-scale metrics differed significantly in their responses to drivers (*QM_3_* = 11.356, *p* = 0.01). Furthermore, a mixed-effects meta-analysis including the interaction term ‘taxon × *⍺*-scale metric’ yielded lower AIC values than the reduced additive model, indicating that specific taxa respond differently depending on the metric measured. Consequently, the breakdowns of mean effect sizes for specific combinations are provided in the Supporting Information (SI).

The meta-analyses summarizing effects on *β*-scale metrics of microbial communities clearly showed strong shifts in community composition in response to global change drivers (Fig. 2, 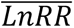 =0.502; 95 % *CI* = 0.416 to 0.588; *p* < 0.0001, based on *n* = 2,433 effect sizes; insufficient data precluded further stratification of the drivers.). Magnitude of community shifts strongly differed between drivers (*QM_4_* = 14.702, *p* = 0.005, albeit a Multiple Comparisons of Means yielded no significant Tukey Contrasts).

Global biodiversity and ecosystem functions are increasingly threatened by anthropogenic activities (Brondízio et al. 2019). Effective and resource-efficient nature conservation therefore requires detailed knowledge of how global change drivers differentially affect organismal groups (Keck et al. 2025). In this context, ranking the drivers that influence microbial diversity, functioning, and community composition is essential for identifying the most critical pressures, thereby providing the basis for prioritizing microbe-targeted conservation strategies. Land-use and land-cover change and specific pollutants emerged as the dominant drivers of microbial decline (Figs. 1 and 2), consistent with findings for macroscopic taxa (Keck et al. 2025). Although many drivers exhibited mean negative effects on microbial α-scale metrics the frequent overlap of their 95% confidence intervals with zero indicates a strong heterogeneity and potential context dependency of microbial responses. Such variability likely arises from heterogeneous exposure to perturbations at the micro-scale, as well as from complex interactions among microbial taxa and their abiotic environment that modulate these responses (Lee et al. 2025). Context dependence also arises from the synchronous occurrence of multiple perturbations and stresses. Two or more global change drivers may have additive, synergistic, or antagonistic effects that are often not predictable from the responses to each driver in isolation (Rodríguez Del Río et al. 2025; Zhou et al. 2020). Accordingly, the combined effects of multiple global change drivers should be more systematically considered when assessing threats to microbial diversity and function. Furthermore, meta-analytical coverage of microbial functionality remains fragmented. Consequently, the SOMA presented here primarily captures broad biogeochemical process rates, such as soil respiration and nitrification. This focus likely overlooks critical shifts in specific functional genes. Crucially, recent studies demonstrate that global change drivers can select for detrimental traits, such as pathogenicity and antibiotic resistance, which are accumulating in the environment (Rodríguez Del Río et al. 2025; Geng et al. 2025). This often undetected functional shift underscores the urgent need for comprehensive metagenomic monitoring strategies to detect and track these emerging threats. Crucially, long-term monitoring should be implemented for establishing urgently needed baseline data and for tracking temporal changes in taxonomic and functional diversity and composition, as declines may accumulate gradually and only become apparent once critical thresholds are reached (Sun et al. 2025).

### Instrumental rationales

For the second SOMA, I conducted a Web of Science search for meta-analyses addressing microbial (bacteria, fungi, the total microbial assemblage) roles in ecosystem functions, element cycles, and host performance (methods see above and SI). Synthesizing 6,452 effect sizes, I found that individual microbial strains or microbial diversity improves ecosystem functions and host performance by an average of 35.9% (estimated weighted mean effect size 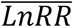 = 0.307, 95 % confidence interval (*CI*) = 0.214 to 0.400, *p* < 0.0001). Specifically, there is robust evidence for positive microbial effects on critical processes including organic matter decomposition, nutrient cycling (C and N), and plant growth, as well as disease suppression and the reduction of heavy metal accumulation in plants (Fig. 3A). This general trend is corroborated by the consistently positive (and frequently significant) mean effect sizes observed across the individual meta-analyses underlying this synthesis (Fig. 3A). The effect sizes differed between microbial functions differed (multivariate meta-analysis model, Q-test for moderators: *QM_4_* = 25.13, *p* < 0.0001), but not between microbial taxa (bacteria or fungi, *QM_2_* = 0.82, *p* = 0.663).

The results presented here clearly confirm the instrumental value of microbes in ecosystems. The strong and consistent positive effects on ecosystem functions and host health as shown in the first SOMA align with essential functions ranging from microscale interactions to global biogeochemical cycles. For instance, at small scales, microbes enhance the fitness of their eukaryotic hosts (Trevelline et al. 2019; Afkhami et al. 2025), while at global scales, the absence of prokaryotic organisms would lead to nitrogen accumulation in the oceans and a depletion of fixed nitrogen on land (Gilbert and Neufeld 2014). Beyond present-day functions, microbe-mediated contributions are instrumental for safeguarding the planet’s future. Microbes are increasingly recognized as essential agents for enabling hosts and ecosystems to acclimate to global change (Peixoto et al. 2022; Baldrian et al. 2023; Zieschank et al. 2025), as well as for achieving many of the United Nations Sustainable Development Goals (Borgianni et al. 2025). Beyond these ecological functions, the metabolic capabilities of microbes offer potential tools for technological solutions to mitigate climate change (Peixoto et al. 2024) and improve human and environmental health within a One Health framework (Peixoto et al. 2022; Banerjee and Van Der Heijden 2023). Furthermore, a strong argument for maintaining microbial diversity lies in its option value arising from as-yet-unknown functions and evolutionary innovations that may become essential for future ecosystem adaptation or biotechnological solutions (Han et al. 2023). Consequently, the precautionary principle implies a need for microbial conservation to prevent the loss of this future potential. Finally, while potential negative effects such as pathogenicity must be considered, the risks associated with conserving natural, healthy microbiomes – where pathogens often fulfill specific ecological roles (Ingram 2022) – are generally considered lower than the risks posed by active interventions such as artificial inoculations (Peixoto et al. 2022).

### Existing conservation interventions

For the third SOMA, I conducted a Web of Science search for meta-analyses summarizing evidence of effects of existing conservation interventions such as afforestation, organic agricultural practices, or restoration on microbial α-scale metrics (methods see above and SI). Overall, existing conservation interventions positively affected *⍺*-scale metrics of microbial communities (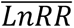 =0.4575; 95 % *CI* = 0.155 to 0.760; *p* < 0.003, based on *n* = 4,058 effect sizes) with no differences between intervention types (*QM_2_* = 0.312, *p* = 0.856). However, both *⍺*-scale metrics (abundance, biomass, diversity, or function; *QM_3_* = 43.198, *p* < 0.0001) and taxa (bacteria, fungi, or the total microbial assemblage; *QM_2_* = 25.135, *p* < 0.0001) differed strongly in conservation gain. Only microbial biomass significantly recovered due to conservation interventions, whereas microbial diversity, abundance, and functions responded highly heterogeneously (Fig. 3B). *⍺*-scale metrics of fungi and total microbial assemblages showed gains from conservation interventions, whereas bacteria exhibited no consistent response (Fig. 3B). This heterogeneity is further supported by the low proportion of significantly positive effect sizes reported in the underlying meta-analyses, a pattern particularly evident for diversity metrics and, to a lesser extent, functionality across both fungal and bacterial communities (Fig. 3B, pie charts). A mixed-effects meta-analysis including the interaction term taxon × *⍺*-scale metrics yielded a slightly higher AIC value than the reduced additive model, confirming no significant interaction. For completeness, however, the specific mean effect sizes for different taxon-metric combinations are provided in the Supporting Information (SI).

Nature conservation interventions, such as the protection and sustainable management of areas, reduction of habitat loss, and restoration, have been shown to substantially improve the state of plant and animal biodiversity (Langhammer et al. 2024). In contrast, such interventions appear less consistently effective for microbial recovery. Success seems highly dependent on microbial taxa and *⍺*-scale metrics, and, given the high variability observed even within specific taxon-intervention combinations, most likely on the environmental context (Fig. 3B, SI). For instance, ecological restoration often has unpredictable effects on microbial communities, and even if they recover in diversity, microbial functionality may not reach the levels found in natural ecosystems (Hart et al. 2020). Restoration and other interventions may be enhanced by adding measures that specifically address microbes, which supports microbial recovery and thereby also ecosystem processes and functions at the macroscopic level (Contos et al. 2025). Microbe-targeted conservation interventions may include measures ranging from enhancing local microbial reservoirs – such as leaf litter, dead wood, and field margins – to microbial transplants or inoculations. However, particularly the latter approaches may carry risks, including the spread of pathogens and the disruption of existing microbial interaction networks (Peixoto et al. 2022). While it has been argued that these risks are controllable and outweighed by the costs of inaction (Peixoto et al. 2022), conserving natural ecosystems and increasing the number and types of local reservoirs of microbial diversity may represent the most promising and effective means of maintaining microbial functions (Contos et al. 2025; Hemraj et al. 2024).

### Vulnerability and conservation gain

The concept of vulnerability is a cornerstone of conservation ecology, as it enables the identification of the most threatened ecosystems and organisms and supports the effective allocation of limited conservation resources (Fig. 4A). Vulnerability is composed of three components: exposure, sensitivity and adaptive capacity (Turner et al. 2003; Foden et al. 2013). The SOMA on threats confirms that microbes are both exposed and sensitive to specific global change drivers (Shade et al. 2012). However, the magnitude of this susceptibility likely varies across organizational levels: individual cells are exposed when directly encountering a stressor, whereas populations and communities can be buffered against global change drivers by spatial heterogeneity, which modulates the intensity of exposure. Different organizational levels also rely on distinct mechanisms to adapt to global change drivers. Individual cells utilize molecular protection systems, such as heat shock proteins, antioxidant defenses, and DNA repair mechanisms (Guan et al. 2017). At the population level, microbes utilize rapid growth, short generation times, and high mutation rates, alongside horizontal gene transfer, to facilitate rapid evolutionary adaptation (Shade et al. 2012). Communities, in turn, may exhibit resistance (remaining unchanged) or resilience (quickly recovering original diversity, composition, and functionality). Alternatively, functional redundancy among community members allows for functional homeostasis even when community composition changes (Shade et al. 2012). Accordingly, the pronounced shifts in community composition observed under global change drivers (Fig. 2) may reflect either a sensitivity to these drivers (a collapse) or an adaptive response (species turnover) that maintains functional stability. Future research should integrate assessments of exposure, sensitivity, and adaptive capacity across organizational levels to establish a robust framework for microbial vulnerability.

**Fig. 4.**
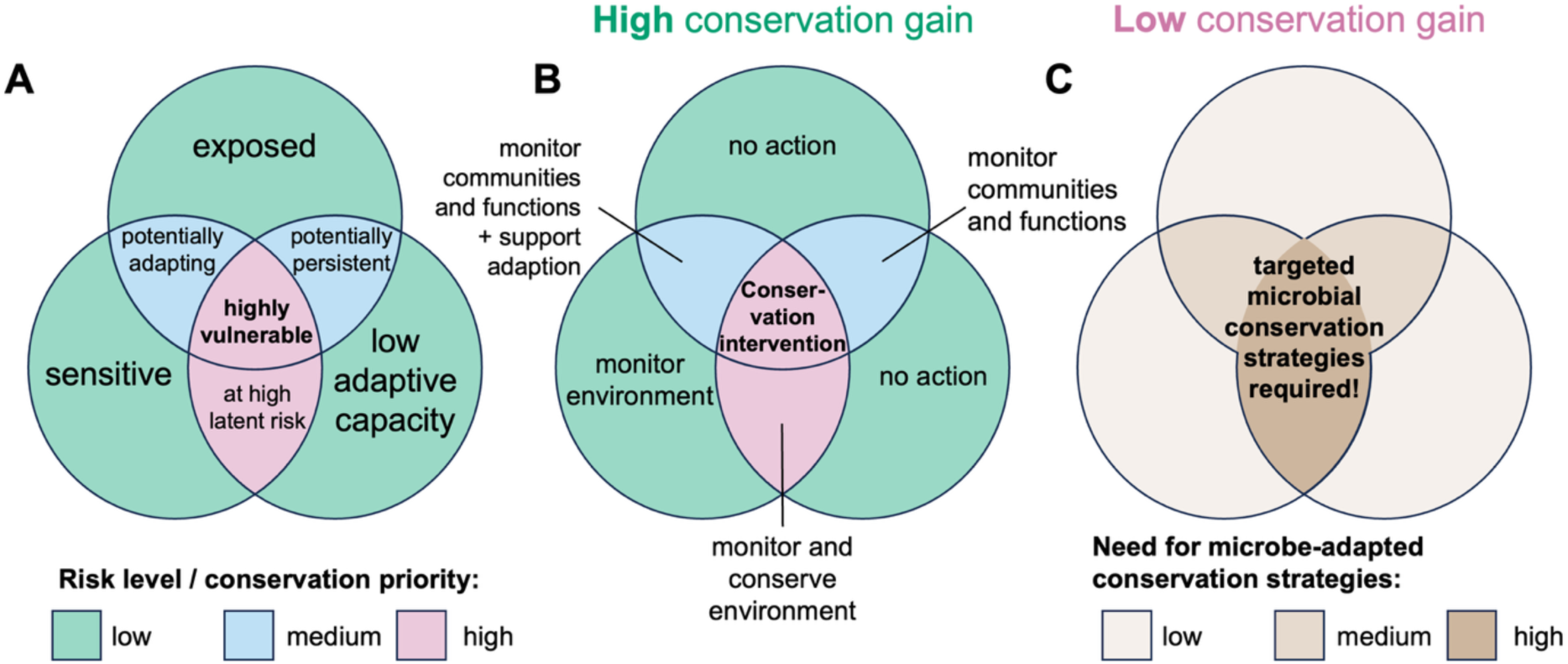
Vulnerability and conservation gain. Specific combinations of exposure, sensitivity adaptive capacity indicate low (green), medium (blue), or high (pink) risk levels and resulting conservation priorities (**A**), along with recommendations for action (Foden et al. 2013) (**B**). The action recommendations, however, are meaningful only if the targeted organisms are expected to exhibit a high recovery potential and thus achieve a conservation gain in response to conservation interventions. As existing conservation interventions often fail to protect and restore microbes – indicating a low conservation gain – there is a clear need to design and implement targeted microbial conservation strategies (**C**).

The findings of this study suggest that microbes can be regarded as vulnerable across multiple contexts, warranting conservation prioritization and targeted actions (Fig. 4B) (Foden et al. 2013). However, such recommendations are practically meaningful only if the targeted organisms exhibit a high recovery potential, thereby allowing for a tangible conservation gain in diversity, composition, or function. The International Union for Conservation of Nature (IUCN) addresses this through the Green Status of Species framework, which evaluates the impact of conservation actions and reveals that most plant and animal species respond positively to intervention (Grace et al. 2021). In stark contrast, existing conservation interventions often fail to consistently maintain or restore microbial community diversity, composition, and function (Fig. 3B). Given this discrepancy, distinct and targeted microbial conservation strategies are urgently required to safeguard microbial diversity and habitats, thereby securing ecosystem functioning and host health (Fig. 4C).

## Conclusion

The SOMAs presented here synthesize robust evidence that microbes (like plants and animals) are threatened by anthropogenic activities, underscoring the urgent need for targeted microbial conservation. However, developing these strategies presents a unique challenge: while some existing conservation frameworks may require only minor modifications to account for microbial characteristics, others may prove difficult or impossible to adapt to the distinct biological requirements of the microbial world (Junker and Farwig 2025). The recently passed EU Soil Monitoring Law (Directive (EU) 2025/2360 of the European Parliament and of the Council of 12 November 2025 on Soil Monitoring and Resilience (Soil Monitoring Law) 2025) specifically addresses the monitoring of soil biodiversity including microorganisms, which is an important first step towards microbial conservation. Future efforts should expand to include other microbial habitats and hosts, as well as the development of microbe-targeted conservation interventions to prevent microbiome degradation, preserve natural reservoirs of microbial diversity, and restore degraded sites. The critical first step toward effective strategies is to address existing research gaps and build the robust knowledge base needed to inform policy and practice. Specifically, a mechanistic understanding of the processes shaping the global, regional, and local distribution of microbial diversity—as well as the occurrence patterns of specific taxa and functions—is essential. This knowledge is required to identify priority areas for conservation, encompassing both microbial biodiversity hotspots and the most vulnerable assemblages (Guerra et al. 2022). Crucially, the availability of affordable high-throughput sequencing methods to monitor and geographically map functional genes, alongside taxonomic units, will be instrumental in supporting these goals, particularly as traditional species-level protection remains largely unfeasible for the microbial world (Junker and Farwig 2025). Consequently, the ongoing debate regarding the protection of species versus ecological functions (Dee et al. 2017; Tobias et al. 2025) is effectively obsolete in the microbial context. Instead, priority should be assigned to interventions that safeguard ecosystem functions, particularly within communities characterized by high functional redundancy. Conversely, where microbial taxa are functionally distinct, the goals of species and functional conservation effectively converge. In these instances, the status of functionally unique taxa must be closely monitored and, where necessary, supported through targeted conservation measures (Litchman et al. 2024). Following this knowledge acquisition, the design and implementation of specific interventions constitutes the critical second step toward effective microbial conservation. Promising approaches to maintain and enhance microbial diversity and functioning include microbe-inclusive ecological restoration, the augmentation of habitats with natural reservoirs for microbial diversity, and the targeted protection of host organisms (Trevelline et al. 2019; Contos et al. 2025). Ultimately, saving microbes means protecting the very foundation of life – safeguarding ecosystem integrity, planetary health, and the resilience of all living systems. Consequently, it is imperative that microbes are recognized, studied, and protected as fundamental components of biodiversity conservation and ecosystem management.

## Acknowledgements

I thank Lars Voll for inspiring this project by persistently asking for evidence that global change threatens microbes; Katarzyna Roguz, Luise Szymanski, and Alexander Zizka for constructive feedback on an earlier version of the manuscript; and many colleagues for fruitful discussions on microbial conservation. I am grateful to Hamed Azarbad, Karl-Heinz Rexer, Alexander Lach, Annette Schriever, and Christiane Siebert-Morawietz. Their support allowed me to take the time during my sabbatical to work on this manuscript.

## Author contributions

RRJ designed the study, performed the second-order meta-analyses, and wrote the manuscript.

## Supplementary Information

Supplementary Information: Supplementary methods, supplementary figures and tables, references to original meta-analyses, supplementary results.

## Data Availability

Data are available at figshare.com (https://doi.org/10.6084/m9.figshare.30875873).

## Supplementary Information

### Methods

#### Search terms, PRISMA, and included meta-analyses

I used the following search terms for the three second-order meta-analyses:

<u>Instrumental rationales:</u> TI=(metaanalys* OR “systematic review” OR (meta NEAR/1 analys*)) AND TS = (microbiome OR microbiota OR “microbial communit*” OR “microbial diversity” OR bacteria* OR fung* OR virus* OR protist* OR archaea OR prokaryot* OR micro?algae) AND TS = (“Ecosystem service*” OR “ecosystem function*” OR “plant growth promot*” OR “*degradation of pollutions” OR “host health” OR “host fitness” OR “host performance” OR “beneficial function*” OR “soil formation” OR “nutrient cycling” OR “climate regulat*” OR “disease control” OR “biodiversity-ecosystem function” OR “provisioning service*” OR “regulating service*” OR “cultural service*” OR “supporting service*” “decomposit*” OR “water purification”)

<u>Threat-based rationales:</u> TI=(metaanalys* OR “systematic review” OR (meta NEAR/1 analys*)) AND TS = (microbiome OR microbiota OR “microbial communit*” OR “microbial diversity” OR bacteria* OR fung* OR virus* OR protist* OR archaea OR prokaryot* OR micro?algae) AND TS = (“global change” OR “land-use change” OR “land use” OR “habitat loss” OR “sea use” OR “sea-use change” OR urbanization OR pollution OR “invasive species” OR invasi* OR “alien species” OR “exotic species” OR “climate change” OR “climate warming” OR “global warming” OR “temperature rise” OR drought OR “precipitation change” OR “nitrogen deposition” OR fertilization OR exploitation OR overharvest*)

<u>Effects of existing conservation interventions:</u> TI=(metaanalys* OR “systematic review” OR (meta NEAR/1 analys*)) AND TS=(microbiome OR microbiota OR “microbial communit*” OR “microbial diversity” OR bacteria* OR fung* OR virus* OR protist* OR archaea OR prokaryot* OR micro?algae) AND TS=(“conservation measure*” OR restoration OR rewilding OR rewetting OR “deadwood addition” OR afforestation OR “habitat creation” OR “habitat protection” OR “protected areas” OR “habitat corridor*” OR “ecological corridor*” OR “agroecological structure*” OR “flower strips” OR “field margins” OR hedgerows OR “set-aside” OR regeneration OR “buffer zones” OR “species protection” OR “legal protection” OR “nest site*” OR “low-input farming” OR “organic farming” OR intercropping OR “green infrastructure”)

I screened the title and abstract of each article to assess its eligibility for inclusion in the final analysis. The selection steps and resulting study numbers, including exclusions, are shown in PRISMA flow charts (Fig. SI.1):

**Fig. SI.1.**
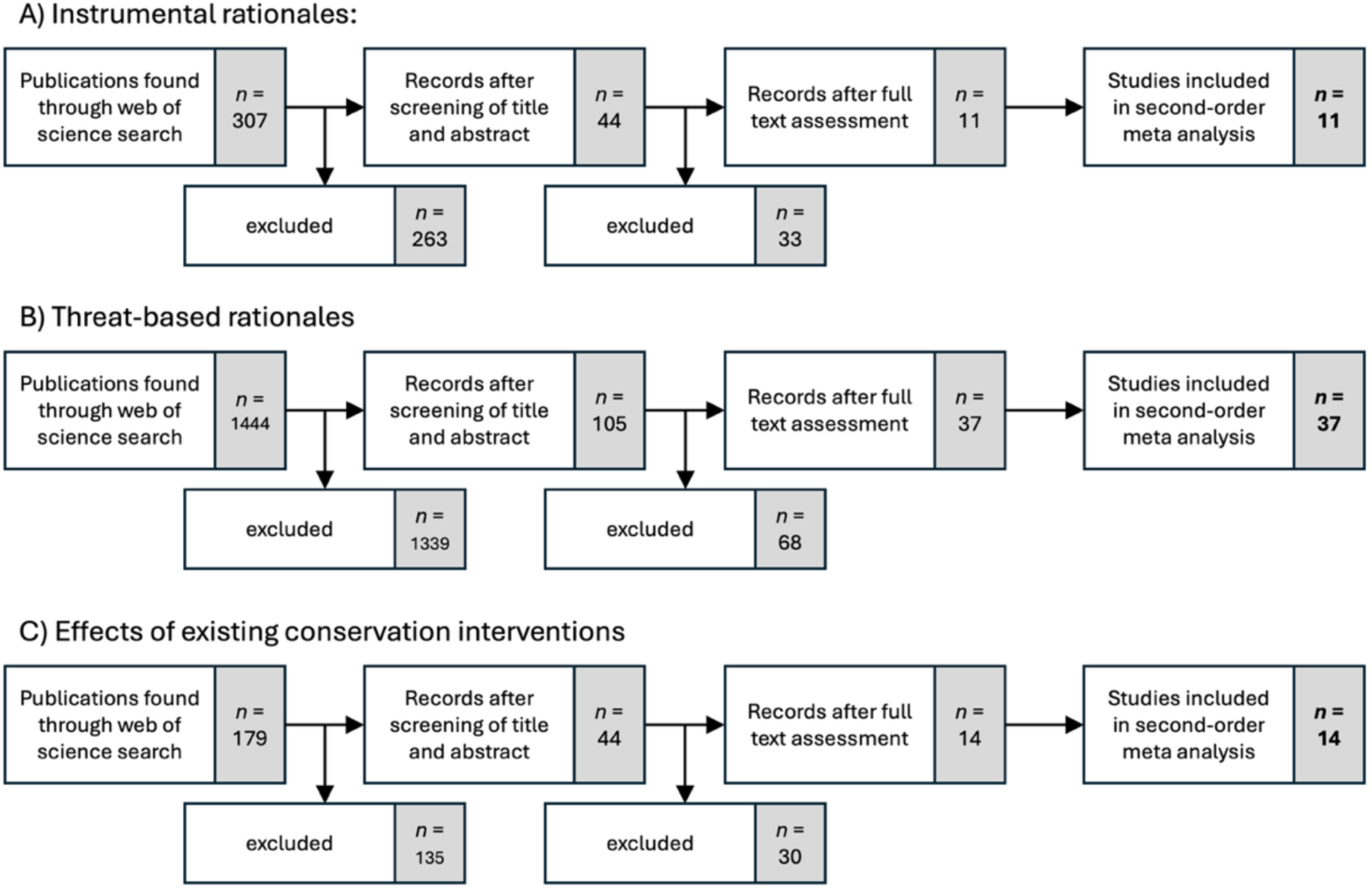
Prisma flow charts.

The following meta-analyses were used for the second-order analyses:

## Supporting results

### Threat-based rationales

Most of the estimates shown in Fig. 2 and 3 of the main text did not indicate meaningful or strong small-study effects or publication bias. For land conversion and N-addition in Fig. 1, Egger’s test yielded intercepts that slightly deviated from zero (*p* = 0.046 or *p* = 0.039, respectively), suggesting a weak small-study effect. However, the funnel plot for land conversion (Fig. SI.2A) exhibited little visible asymmetry; this signal therefore is unlikely to influence the interpretation of the result. The funnel plot for N-addition (Fig. SI.2B), however, shows clear signs of asymmetry and thus results may be interpreted with some caution.

**Fig. SI.2.**
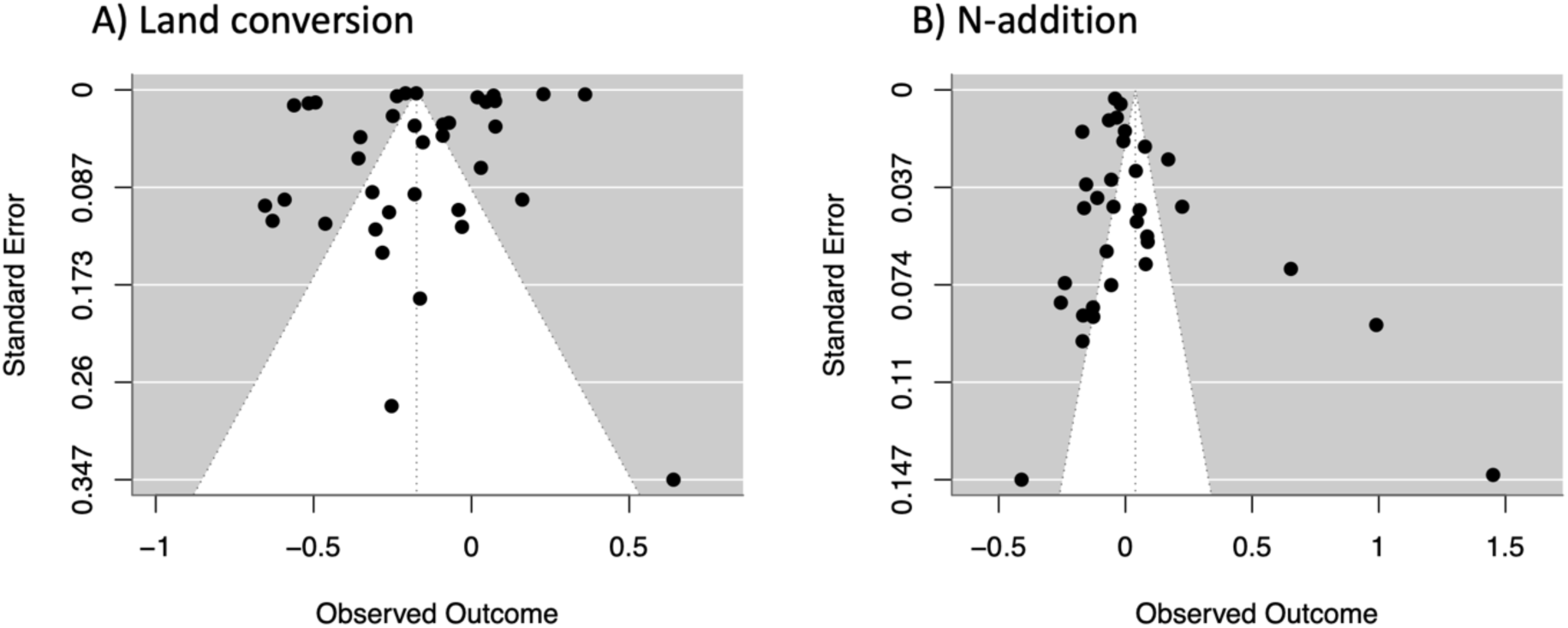
Funnel plot of effect sizes for the effects of A) land conversion and B) N-addition on microbial communities.

While the effect sizes did not differ between microbial taxa (*QM_2_* = 1.690, *p* = 0.430) indicating taxon-independent effects, the types of *⍺*-components (diversity, abundance, biomass, function) did differ in their responses to drivers (*QM_3_* = 11.356, *p* = 0.01). Accordingly, mixed-effects meta-analysis containing the interaction term taxon * *⍺*-component as moderator received lower AIC-values than the reduced model containing additive effects only. To summarize effects of global change drivers on *⍺*-components of different taxa, mean effect sizes of different combination are given in Tab. SI.1, which may be consulted to identify the highest priorities for microbial conservation measures.

### Instrumental rationales

The estimates shown in Fig. 1 of the main text each originate from between one to nine meta-analyses, only, that however consider large numbers of effect sizes. Therefore, I could not meaningfully test for small-study effects or publication bias in most cases, except for microbial effects on plant growth, which showed no signs of a small-study effect (Egger’s test: *p* = 0.0782).

### Effects of existing conservation interventions

Most of the estimates shown in Fig. 4 of the main text did not indicate meaningful or strong small-study effects or publication bias. For microbes and function in Fig. 4, Egger’s test yielded intercepts that deviated from zero (*p* = 0.0107 or *p* = 0.0464, respectively), suggesting small-study effects and may therefore be interpreted with some caution.

Mixed-effects meta-analysis containing the interaction term taxon * α-component as moderator received slightly higher AIC-value than the reduced model containing additive effects only indicating no significant interaction term. Nonetheless, to summarize effects of conservation measures on α-components of different taxa, mean effect sizes of different combination are given in Tab. SI.2.

**Tab. SI.1.**
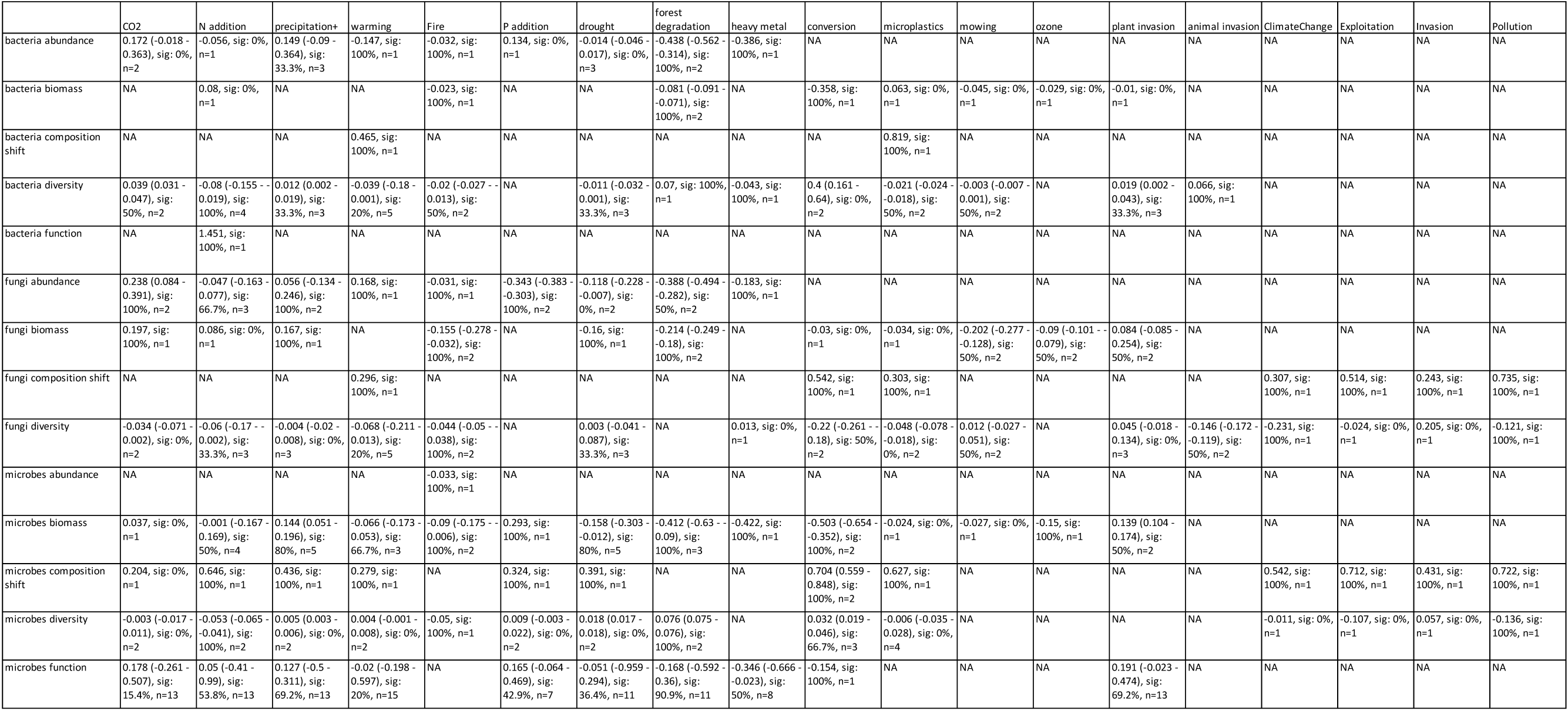
Summary of meta-analyses on threat-based rationales. Mean effect sizes of taxon *α-component combinations are given. Additionally, the range of effect sizes in original meta-analyses, the percentage of significant effects sizes, and the number n of meta-analysis reporting each combination is given.

**Tab. SI.2.** Summary of meta-analyses the effects of existing conservation interventions. Mean effect sizes of taxon * α-component combinations are given. Additionally, the range of effect sizes in original meta-analyses, the percentage of significant effects sizes, and the number n of meta-analysis reporting each combination is given.

|  | intercropping | crop rotation | no fertilizer | no tillage | organic fertilizer | reduced tillage | covercrops | interplanting | organic farming | afforestation | rehabilitation | restoration | no-longer-management | agroforestry | revegetation |
| --- | --- | --- | --- | --- | --- | --- | --- | --- | --- | --- | --- | --- | --- | --- | --- |
| bacteria abundance | 0.179, sig: 0%, n=1 | NA | NA | NA | NA | NA | NA | NA | NA | NA | NA | NA | NA | NA | NA |
| bacteria biomass | 0.228, sig: 0%, n=1 | 0.271, sig: 0%, n=1 | -0.081, sig: 0%, n=1 | -0.045, sig: 0%, n=1 | 0.537, sig: 100%, n=1 | 0.652, sig: 100%, n=1 | NA | NA | NA | NA | NA | NA | NA | NA | NA |
| bacteria diversity | 0.08 (0.035 - 0.126), sig: 50%, n=2 | NA | NA | NA | NA | NA | NA | NA | NA | 0.029, sig: 0%, n=1 | -0.114 (-0.231 - 0.004), sig: 50%, n=2 | 0.024 (-0.013 - 0.08), sig: 20%, n=5 | NA | NA | NA |
| fungi abundance | 0.25, sig: 100%, n=1 | 1.146, sig: 0%, n=1 | NA | NA | NA | NA | 1.343, sig: 100%, n=1 | 1.198, sig: 100%, n=1 | NA | NA | NA | 4.244, sig: 100%, n=1 | NA | NA | NA |
| fungi biomass | 0.379, sig: 100%, n=1 | 0.45, sig: 100%, n=1 | -0.146, sig: 0%, n=1 | 0.249, sig: 100%, n=1 | 0.456, sig: 100%, n=1 | 0.657, sig: 100%, n=1 | NA | NA | 0.635 (0.377 - 0.893), sig: 100%, n=2 | NA | NA | NA | NA | NA | NA |
| fungi diversity | 0.108 (0.099 - 0.116), sig: 0%, n=2 | 1.164 (0.972 - 1.355), sig: 0%, n=2 | NA | NA | NA | NA | 1.119 (1 - 1.238), sig: 0%, n=2 | 1.018 (1.007 - 1.029), sig: 0%, n=2 | -0.031 (-0.17 - 0.025), sig: 25%, n=4 | 0.037, sig: 100%, n=1 | 0.041 (0.038 - 0.044), sig: 50%, n=2 | 0.074 (0.015 - 0.193), sig: 20%, n=5 | 0.167, sig: 100%, n=1 | NA | NA |
| microbes biomass | 1.514, sig: 100%, n=1 | NA | NA | NA | NA | NA | NA | NA | 0.338, sig: 100%, n=1 | 0.482, sig: 100%, n=1 | NA | 0.566 (0.417 - 0.715), sig: 100%, n=2 | NA | 0.225, sig: 100%, n=1 | NA |
| microbes diversity | NA | NA | NA | NA | NA | NA | NA | NA | NA | NA | NA | NA | NA | 0.086, sig: 100%, n=1 | 0.075 (0.051 - 0.099), sig: 50%, n=2 |
| microbes function | 3.079 (2.347 - 3.812), sig: 100%, n=2 | NA | NA | NA | NA | NA | NA | NA | 0.335 (-0.102 - 0.613), sig: 75%, n=4 | 0.624 (0.177 - 1.094), sig: 83.3%, n=6 | NA | 0.407 (-0.028 - 0.704), sig: 83.3%, n=6 | NA | 0.229 (0.063 - 0.349), sig: 75%, n=4 | NA |

